# Interpretable Decoding of Frequency-Resolved Functional Connectivity

**DOI:** 10.64898/2026.08.20.745932

**Authors:** Santeri Ruuskanen, Eero Saarro, Carola Maria Caivano, Lauri Parkkonen, Ivan Zubarev

## Abstract

Whole-brain functional connectivity, estimated from magnetoencephalography (MEG) data, provides a compact representation of long-range neuronal communication, making it suitable for predictive biomarker discovery. In this work, we propose a deep learning framework (FC-CNN) for predicting brain states from frequency-resolved functional connectivity estimates derived from resting-state MEG recordings. We systematically compare the performance of FC-CNN to that of conventional regression methods using amplitude and phase-based functional connectivity in the well-studied age-prediction task on the Cam-CAN cohort (*n* = 576). We show that FC-CNN outperforms conventional approaches, and that, compared to phase synchronization, amplitude envelope correlation consistently leads to higher prediction performance. Moreover, we present quantitative evidence that the weights of a trained deep learning model can enable neurophysiological interpretation of the activity patterns that inform successful predictions. Our work demonstrates that the proposed approach successfully decodes brain states from MEG functional connectivity and is promising for discovery of predictive biomarkers for brain disorders.

## 1 Introduction

The growing availability of large-scale standardized neuroimaging datasets and sophisticated pattern recognition approaches provides exceptional opportunities to develop new models of brain structure and function across the lifespan. The goal of these efforts is to develop robust and interpretable biomarkers for risk assessment, early detection, diagnosis, phenotyping, and prognosis for a variety of neurological and psychiatric conditions [1]. The development of such predictive models has the potential to not only provide diagnostic tools but also to uncover the underlying neural patterns and inform the development of targeted therapeutic approaches.

Most efforts to develop neuroimaging biomarkers have so far relied on structural, metabolic, and haemodynamic imaging (magnetic resonance imaging (MRI), positron emission tomography, and functional MRI, respectively) [2–9]. A growing number of studies have explored the use of direct non-invasive measurements of electrical brain signaling, including magnetoencephalography (MEG) and electroencephalography (EEG) [10–17]. MEG, particularly when combined with structural MRI and source modeling, has the advantage of combining neural-scale time resolution with precise anatomical information [18] to estimate functional connectivity across large-scale cortical networks [19–21].

Functional connectivity characterizes statistical dependencies between cortical oscillations at different brain regions across a range of physiological frequencies and reflects inter-areal communication in large-scale neuronal networks [21, 22]. Often, functional connectivity is estimated from MEG data recorded at rest, allowing for straightforward measurements of healthy and clinical populations. Therefore, MEG or EEG resting-state functional connectivity (rs-FC) has potential as a standardized data representation for large-scale predictive biomarker discovery across multiple datasets. Several studies have reported changes in rs-FC in neurodevelopment [23–25], healthy aging [26–31], and pathologies such as Alzheimer’s disease [32–35], Parkinson’s disease [36–39], and schizophrenia [13, 40]. Functional connectivity is a particularly efficient representation of resting-state MEG measurements, as it summarizes key features of long-range neural communication dynamics over time. It is, therefore, a suitable target for predictive inference algorithms aiming at developing disease biomarkers. So far, efforts to develop MEG-based biomarkers have focused on evoked responses [14], power spectral features [10, 11, 41], and time-domain features [42]. A few studies have explored the use of various functional connectivity approaches [13, 43, 44]. Since rs-FC contains finer-grained information compared to, for example, spectral power [30], it could prove to be a viable component of functional biomarker discovery.

Brain age prediction is a well-defined task with easily available large-scale MEG datasets [45–50], making it well-suited for developing and benchmarking decoding methods. Healthy aging is characterized by widespread changes in brain structure and function, including white and gray matter atrophy, particularly in subcortical structures [51, 52], changes in structural and functional connectivity derived from MRI [53], altered temporal dynamics at the neuronal time scale [46, 54, 55], and changes in MEG functional connectivity [26, 28–31]. However, these changes incur substantial interindividual variability, which has prompted interest in brain age and methods to estimate it from neuroimaging data [56]. The difference between the predicted brain age and chronological age has been termed the brain age delta, which has been suggested as an index of normative aging and could be indicative of an increased risk of cognitive decline and disease [56–60]. Prior work on brain age prediction has successfully utilized fMRI, structural MRI, and MEG features in both unimodal and multimodal models [61–67]. Several studies have demonstrated the feasibility of brain age prediction from MEG rs-FC, showing that including MEG features improves prediction performance compared to MRI data alone [61, 63, 64]. However, these investigations have relied on general-purpose regression methods instead of algorithms tailored for decoding rs-FC.

Deep neural networks (DNNs) [68] have demonstrated significant advantages in predictive power over traditional machine learning methods in many domains. Yet, most neuroimaging studies rely primarily on simpler linear techniques combined with feature extraction due to the lack of interpretability techniques for more sophisticated models [69]. Prior work on brain age prediction from MEG data has primarily utilized standard machine learning algorithms, and those who have explored deep learning approaches have not attempted to interpret the model activation patterns [63]. Thus, poor interpretability remains a major limiting factor in the wider adoption of DNNs [70]. To better understand which features inform the model predictions, we present a novel pipeline for extracting predictive patterns from the trained DNNs and propose a validation procedure based on ablation analysis.

In this study, we investigate how the rs-FC estimation method, as well as feature extraction and prediction algorithms, affect predictive performance in the brain age prediction task. We introduce a convolutional neural network (FC-CNN) [68] that outperforms conventional predictive regression methods and provides a framework for identifying the most informative nodes. FC-CNN combines feature extraction and prediction in a single algorithm, informed by the spectrospatial structure of the frequency-resolved rs-FC data. The model learns appropriate feature representations from the training data by minimizing the estimation error. We evaluated the performance of this approach using MEG resting-state functional connectivity from 576 individuals in the Cam-CAN database aged 18–87 years (Fig. 1) [45, 71]. Using the identified informative nodes, we performed a model ablation study and found that a model restricted to the most informative nodes performed nearly as well as the full model. We compared the performance of three variants of FC-CNN against conventional feature extraction and predictive regression methods (Fig. 2) using amplitude-based (Amplitude Envelope Correlation; AEC [21]) and phase-based (weighted Phase-Lag Index; wPLI [72]) functional connectivity estimates as inputs. Finally, we investigated whether estimating functional connectivity in an anatomically-informed space improves the performance of prediction algorithms compared to utilizing sensor-level time courses directly.

**Fig. 1.**
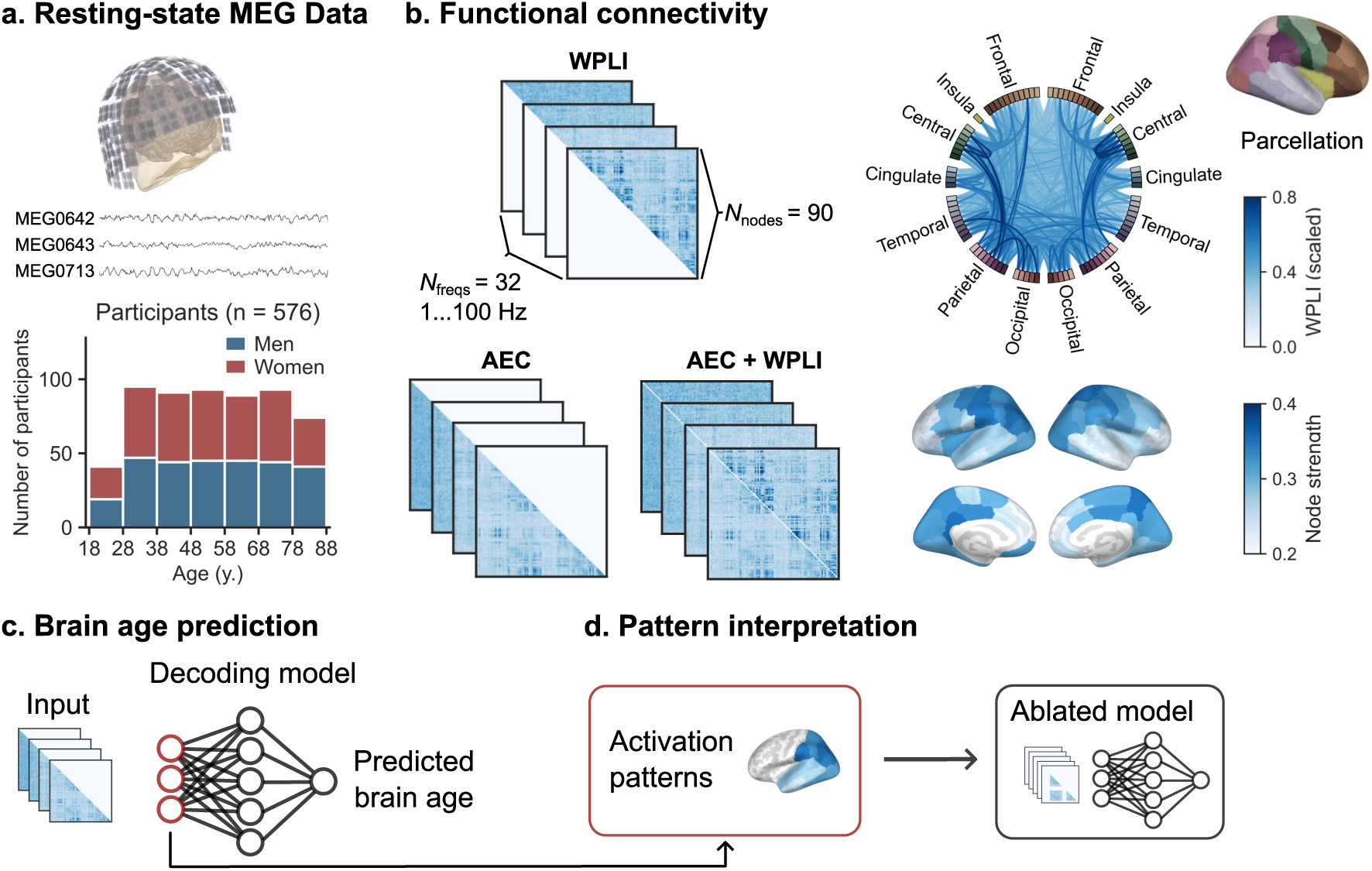
Functional connectivity and age prediction pipeline. **(a.)** We employed MEG data from 576 subjects aged 18–87 years from the Cam-CAN database. **(b.)** We estimated resting-state functional connectivity (rs-FC) using both weighted Phase-Lag Index (wPLI) and Amplitude Envelope Correlation (AEC) between 90 cortical regions of interest (ROIs). **(c.)** Participant age was considered a latent factor, which affects rs-FC through an unknown encoding model. We predicted the brain age through a deep learning-based decoding model. We extracted activation patterns from the weights of the trained decoding model and used the informative features to train an ablated model.

**Fig. 2.**
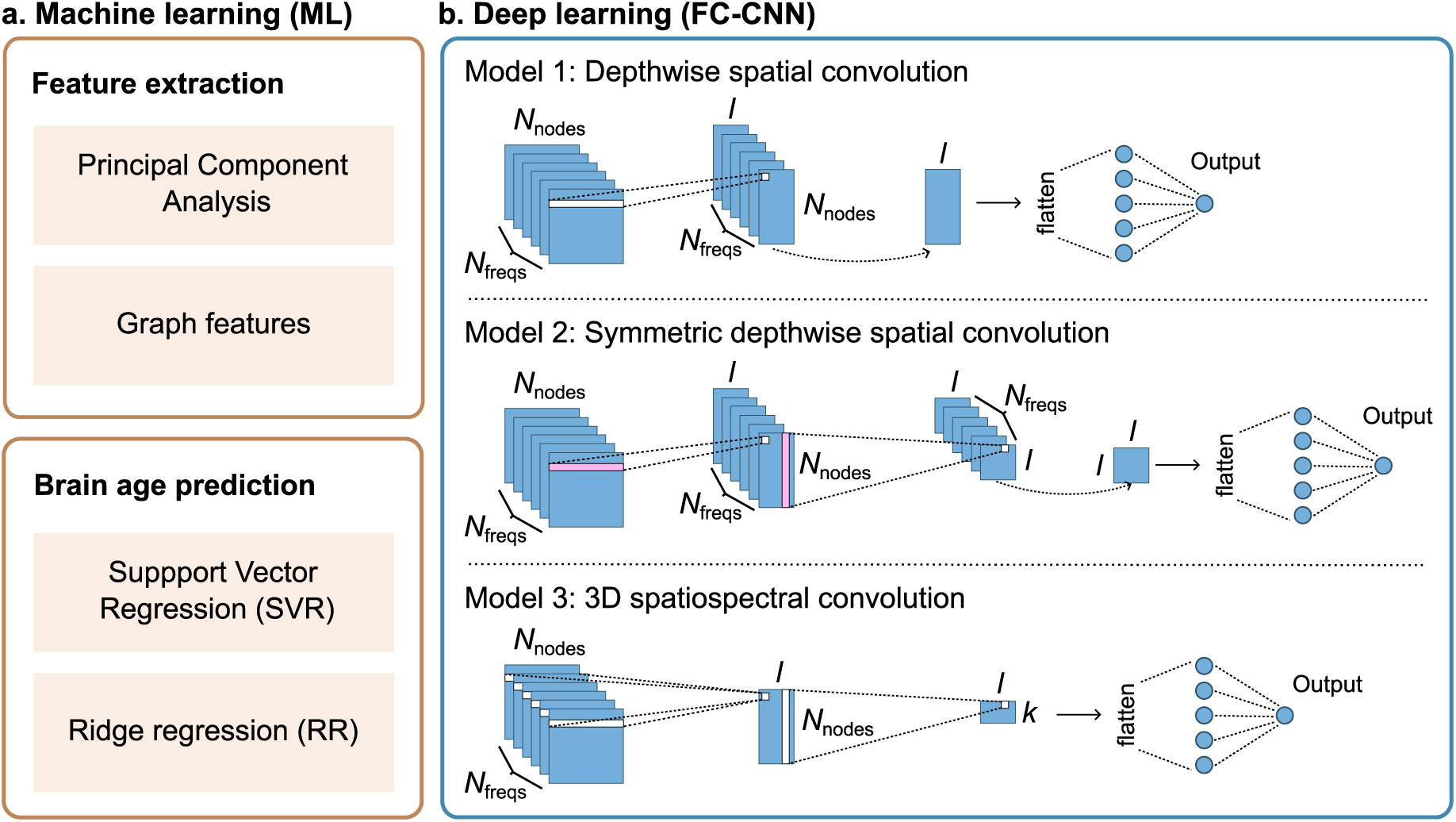
Brain age prediction models. We evaluated two decoding frameworks to predict brain age. The first one consists of feature extraction and age prediction using conventional predictive regression models. The second one combines feature extraction and age prediction in a purpose-built convolutional neural network, FC-CNN. We evaluated three FC-CNN variants with increasing model complexity.

Our work shows that FC-CNN consistently outperforms conventional methods. We demonstrate that exploring the patterns that the model uses to make its predictions can guide efficient feature selection and neurophysiological interpretation of the activity associated with healthy aging. Through model comparison, we found that the FC-CNN variant with the least representational capacity performed best. Our results suggest that amplitude-based rs-FC metrics contain more prediction-relevant information than phase synchronization. We also show that using source-level rs-FC estimates consistently improves the performance of prediction models. These results demonstrate the potential of the FC-CNN architecture and pave the way for interpretable nonlinear decoding methods in functional neuroimaging.

## 2 Results

### 2.1 Deep learning outperformed standard machine learning models

First, we compared the performance of the three proposed deep learning-based decoding models (FC-CNN) to two machine learning models (ML models), namely support vector regression (SVR) and ridge regression (RR), using source-level MEG rs-FC estimates from 576 subjects in the Cam-CAN database. We pooled together the cross-validation (CV) results across all three FC-CNN variants and both ML models trained with rs-FC features derived using Principal Component Analysis (PCA) and graph theory. Each FC-CNN variant was trained three times with different random initializations. Across 12 CV folds and three types of rs-FC datasets (AEC, wPLI, and combined AEC + wPLI), this resulted in 324 FC-CNN fits and 96 ML model fits. The FC-CNN variants delivered an average MAE of 8.57 *±* 0.98 years, and the ML models produced an average MAE of 10.67 *±* 0.93 years (Fig. 3). The difference was statistically significant (Mann–Whitney U-test, *U* = 1975, *p* = 1.28 *×* 10*^−^*^38^). The proposed FC-CNN architecture, therefore, outperformed the reference ML models in the age prediction task.

**Fig. 3.**
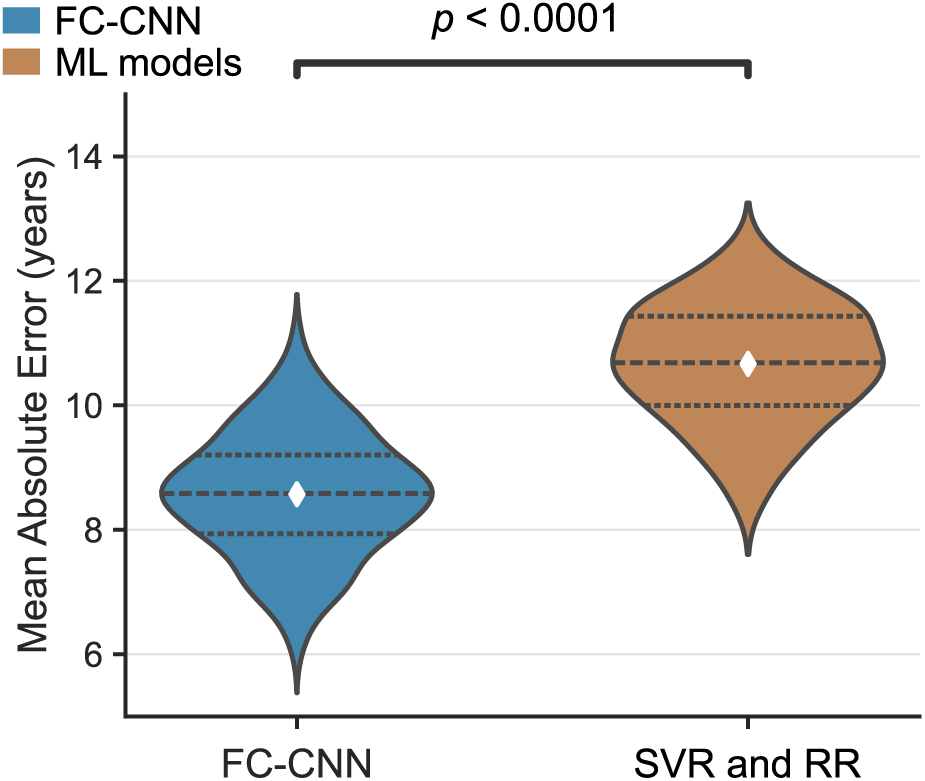
Brain age prediction error of FC-CNN and machine learning models. The prediction errors were aggregated across input datasets, models, folds, runs, and feature extraction techniques. Dashed lines indicate medians, dotted lines indicate quartiles, and white diamonds indicate means. FC-CNN performed statistically significantly better than Support Vector Regression (SVR) and ridge regression (RR) (Mann–Whitney U-test, *U* = 1975, *p* = 1.28 *×* 10*^−^*^38^).

### 2.2 Amplitude envelope correlation is more informative than phase synchronization

Resting-state functional connectivity is typically estimated using either Amplitude Envelope Correlation (AEC; [21]) or phase synchronization metrics such as the weighted Phase-Lag Index (wPLI; [72]). To evaluate the performance of these metrics in the age prediction task, we trained each decoding model using either AEC, wPLI or both as input (Fig. 4). Across the three DL models and pooling together sensor-level and source-level results, wPLI produced an average MAE of 9.63 *±* 1.09 years, AEC provided an average MAE of 8.64.9 *±* 0.97 years, and using both metrics resulted in an average MAE of 8.72 *±* 1.32 years. The difference between wPLI and AEC was statistically significant (Mann– Whitney U-test, two-sided, *U* = 34865, *p* = 3.62 *×* 10*^−^*^18^). However, using both AEC and wPLI did not lead to an increase in model performance. These results suggest that, for the age prediction task, AEC contains a higher degree of relevant information than wPLI.

**Fig. 4.**
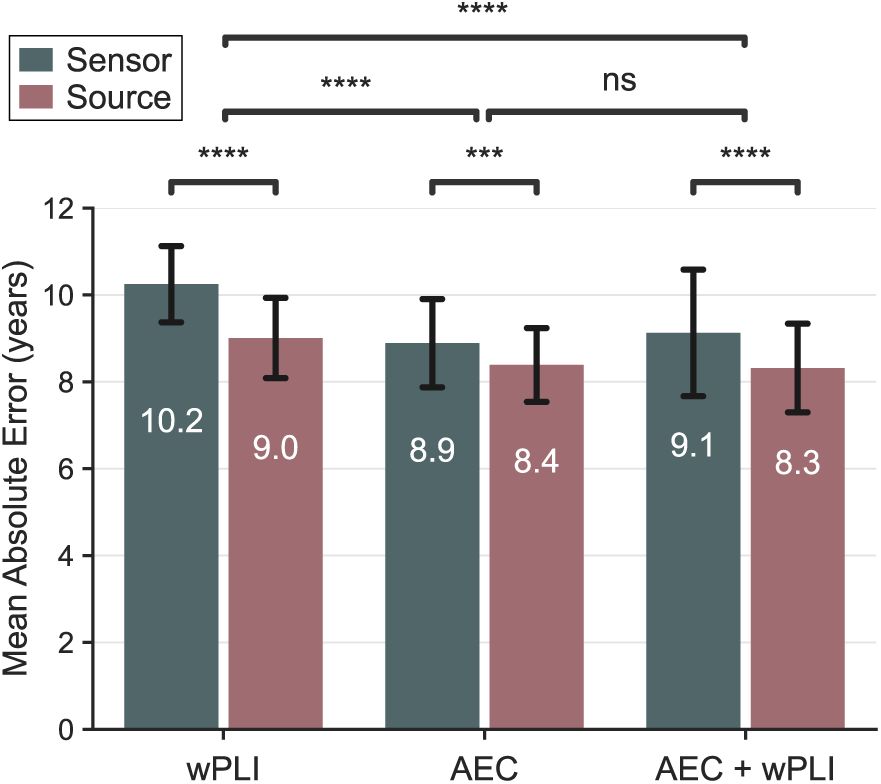
FC-CNN brain age prediction error across input datasets. The results were pooled across the three proposed FC-CNN variants, 12 CV folds, and three runs with different random initializations. The error bars indicate one standard deviation from the mean. The stars indicate statistical significance after Bonferroni correction (Mann–Whitney U-test, two-sided, ***: *p <* 0.001, ****: *p <* 0.0001).

### 2.3 Using an anatomical reference improves prediction accuracy

Frequency-resolved functional connectivity is estimated either from sensor-level signals or time-series that have been projected into an anatomical source space. Although source-space analysis requires additional steps, it is normally preferred to reduce linear mixing between signals from different brain regions [73]. We investigated whether estimating rs-FC in source space is beneficial for age prediction. Using our DL models, we observed significantly higher decoding performance with source-space rs-FC regardless of the functional connectivity metric used (Fig. 4). These findings indicate that functional connectivity should be estimated in an anatomical source space when it is used as input data to a decoding model.

### 2.4 Informative features extracted from the deep learning models

We extracted informative cortical source activation patterns from the weights of the trained deep learning model to identify the cortical ROIs that contributed most to successful predictions using each rs-FC metric. The procedure is described in detail in Methods. Out of the parcellation of 90 ROIs, we identified 31, 26, and 36 informative ROIs for AEC, wPLI, and for combined AEC + wPLI, respectively (Figure 5). These frontal, temporal, and occipital regions largely overlap with regions where we previously observed altered rs-FC [30]. We confirmed the relevance of the resulting patterns for age prediction through model ablation studies, in which a separate model was trained on inputs restricted to the model-derived ROIs (12%, 8% and 16% of input features, respectively). For all connectivity metrics, ablated models performed above chance level explaining 89%, 85%, and 85% of the variation explained by the respective full model.

**Fig. 5.**
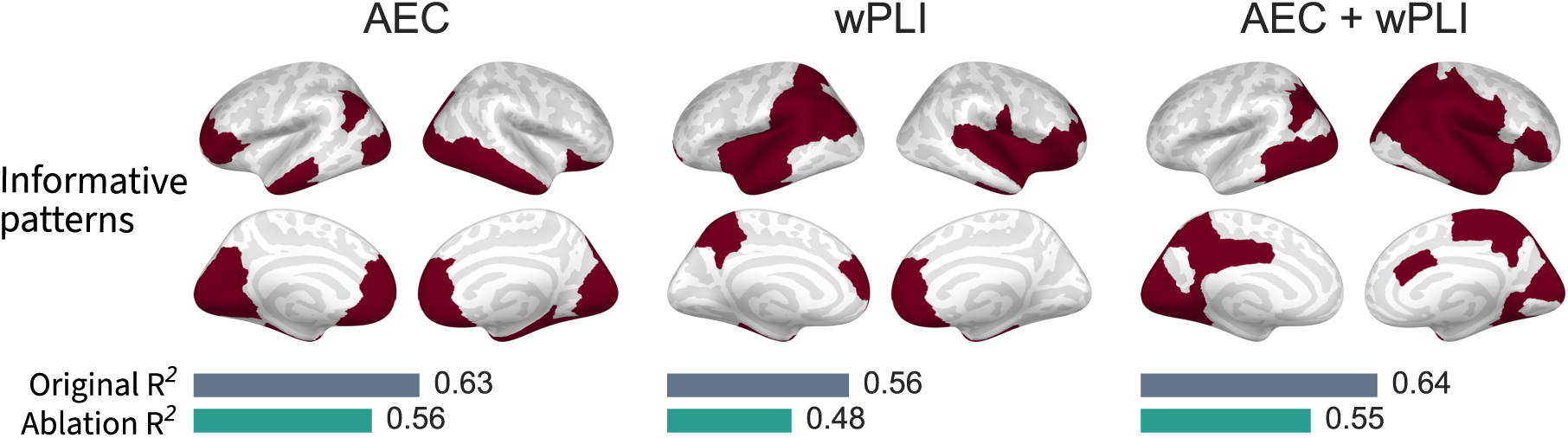
Informative patterns extracted from the weights of the deep learning model. The highlighted cortical regions contributed most to successful brain age prediction in the symmetric depthwise spatial convolution model (Model 2). The bars indicate the coefficient of determination (*R*^2^) of the full model and the ablated model, which utilizes only the informative feature set.

### 2.5 Detailed model comparison

The above discussion considered results aggregated across the three proposed FC-CNN variants and two standard machine learning approaches with PCA and graph-theoretical features. In total, we compared 34 combinations of decoding models and rs-FC features. The decoding performance of each combination is depicted in Fig. 6. In general, the differences between individual models were small compared to the variability across CV folds and model runs. The depthwise spatial convolution model (Model 1) showed the best performance with an MAE of 8.16 *±* 0.92 years. The increased representational capacity of the 3D spatiospectral convolution model (Model 3), did not lead to an improvement in decoding performance. All models performed better than the chance level, an MAE of 15.6 *±* 0.98 years, which was obtained from a dummy model that always predicts the mean age of the dataset in each fold.

**Fig. 6.**
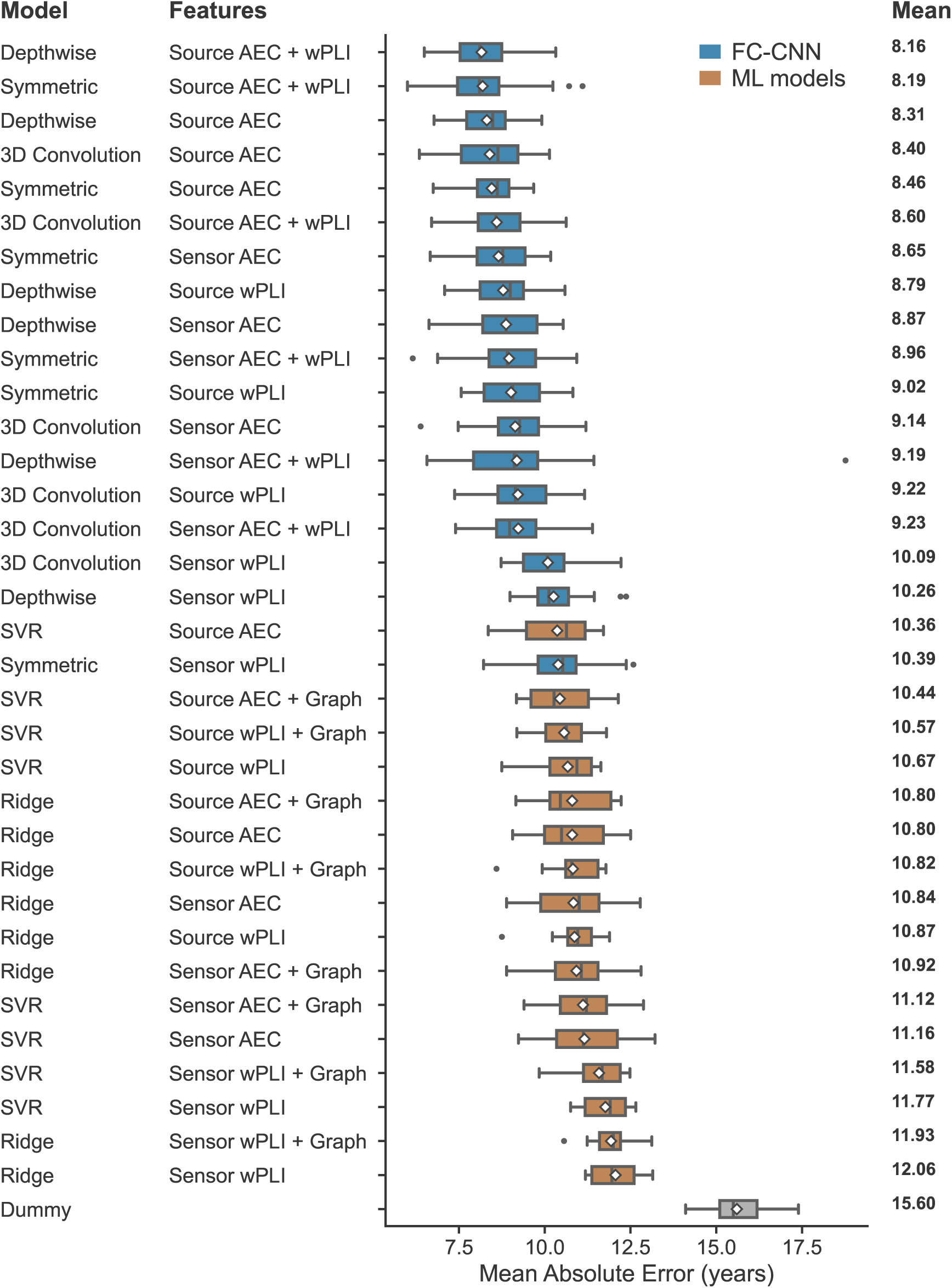
Brain age prediction errors of individual models. The boxes indicate the interquartile range (IQR), the whiskers indicate 1.5 *×* IQR below and above Q1 and Q3, respectively. The vertical black lines indicate the median across cross validation folds and DL model runs. The model–feature combinations are ordered by the mean values (white diamonds).

In contrast to the automatic feature extraction incorporated in FC-CNN, feature engineering and selection is typically required for standard ML models to optimize their ability to capture meaningful patterns in the input data. To capture graph-level information, we combined dimensionality reduction using PCA with graph-theoretical measures, including global efficiency, characteristic path length, and transitivity. The results indicate that incorporating graph-level features generally improved the predictive accuracy (Fig. 6). Nevertheless, SVR and RR still performed worse than FC-CNN.

## 3 Discussion

In this study, we proposed an interpretable deep learning framework for decoding brain states from resting-state functional connectivity (rs-FC) estimated from MEG or EEG recordings. Through extensive comparisons of input features, feature extraction approaches, and predictive model architectures, we demonstrate that: 1) Amplitude Envelope Correlation (AEC) is more informative for brain age prediction than a phase-synchronization metric (wPLI); 2) Convolutional neural networks (FC-CNN) significantly outperform unsupervised feature extraction combined with standard regression models; 3) Anatomically-informed source estimation provides clear benefits to decoding analysis even when deep learning is used; and that 4) informative features can be extracted from a trained deep learning model, increasing model interpretability.

### 3.1 Amplitude-based connectivity is more informative for age prediction

Although previous results suggest that amplitude- and phase-based connectivity estimates may contain non-redundant information [74], understanding how these different metrics contribute to predictive performance remains limited. In our study, using AEC features resulted in significantly higher performance 4 compared to wPLI regardless of the feature extraction method and predictive model type. This result is noteworthy, given that prior work has reported age-related changes primarily in phase-based metrics [26, 28–31]. In our previous study, we reported more widespread age-related changes in wPLI compared to AEC [30]. Moreover, others have suggested that the resting-state AEC connectome is remains unchanged in healthy aging [75]. In light of these results, it is surprising that AEC appears to provide improved performance as a predictive feature. One possible explanation for this discrepancy relates to the reliability and signal-to-noise ratio of the AEC and wPLI methods. Prior work has suggested that amplitude-based metrics have higher test–retest reliability than phase-based measures [76–78], which could explain their more robust predictive performance. On the other hand, the amplitudes and peak frequencies of brain oscillations also change in healthy aging [46, 54, 62, 79]. The fact that AEC retains this amplitude information, while it is ignored by wPLI, could largely explain the elevated predictive performance. In the same vein, prior work suggests that local spectral power features alone can provide predictive performance almost as good as shown here [62]. Furthermore, evidence from earlier studies indicates that AEC and wPLI rs-FC patterns may reflect at least partly separate neuronal mechanisms [74]. To that end, we combined AEC and wPLI inputs in shared rs-FC datasets. This resulted in negligible changes in predictive performance, suggesting that predictions were primarily driven by modulations in amplitude-based rs-FC.

### 3.2 Convolutional neural networks outperform manual feature extraction techniques

Resting state functional connectivity (rs-FC) is a compact and efficient representation of the multi-dimensional MEG time series. Nevertheless, rs-FC data are still very high-dimensional, with only a tiny subset of features containing information relevant for prediction. Thus, extracting informative features is of critical importance. In contrast to unsupervised feature extraction used in combination with SVR and RR, CNNs used in this study learned data representations that minimize the overall prediction error [80]. The superior performance of CNN-based models is likely due to this *supervised* nature of feature extraction.

Among the proposed models, the CNN with the least trainable parameters (depthwise convolution model (Model 1) achieved the best performance (MAE 8.16 *±* 0.92 years), outperforming or matching the state-of-the-art brain age prediction using MEG rs-FC [61, 63, 64], power-spectral features [62], Riemannian geometry embeddings [81], or temporal autocorrelation and other time-series features [67]. This demonstrates the potential of rs-FC as a method to summarize complex MEG data, and the effectiveness of the proposed FC-CNN framework as a decoding tool. However, the age prediction errors observed in these studies utilizing recordings of ongoing neuronal activity, with the respective MAEs in the 8–10-year range, are still notably larger than the approximately 3–5 years achieved using anatomical MRI [65, 66, 82] and multimodal approaches [83]. This discrepancy between structure and function could indicate that alterations in brain structure follow chronological age relatively strictly, whereas brain function affords greater flexibility in healthy aging. Whether such flexibility might reflect compensatory mechanisms in healthy aging remains debatable [84, 85]. Moreover, it has been suggested that although functional imaging measures generally result in higher age prediction errors, they could be sensitive for a broader range of phenotypes than structural imaging-based models and therefore potentially more useful as biomarkers [86].

### 3.3 Anatomically-informed source estimation improves predictive performance

Analyzing rs-FC in an anatomical source space requires obtaining participants’ structural MRIs and complex source modeling steps. Although source space analysis has been shown to reduce spurious connectivity [87], source space analysis *per se* does not provide additional information to the decoder. Thus, in principle, the same information should be available to the decoder in the sensor space. Our results demonstrate that predictive performance improved significantly in the anatomical source space compared to the sensor space.

### 3.4 Informative features can be extracted from a trained CNN

A key limitation of deep learning techniques in the brain imaging domain is their limited interpretability. Here, we employ the interpretable deep neural regression approach to understand activation patterns that inform successful age predictions. This analysis identified that, depending on the rs-FC metric, 8–16% of features accounting for over 80% of variation in age explained by the full feature set. ROIs identified by the model are in line with previous studies. We argue that this approach will benefit the interpretation of neural patterns in other prediction tasks.

### 3.5 Limitations

A common challenge in decoding studies utilizing neuroimaging data is the relatively small number of training samples, which can limit the capabilities of deep learning models [68, 80]. While the dataset applied in our study is large in the MEG context, it falls short of the often millions of samples employed to train large deep learning models. To accommodate for this limitation, we designed the proposed models with moderate representational capacity, minimizing the number of trainable parameters. The limited amount of training samples likely explains why the best results were achieved with the least complex model among those evaluated. Hence, it can be speculated that the model performance could further improve given additional training data. Larger datasets would enable scaling up model complexity, thus capturing more nuanced features in the data. To reduce overfitting on the limited available data, we added Gaussian noise to the input rs-FC, which resulted in a notable performance improvement. Gaussian noise works as a data augmentation and regularization technique by introducing randomness into the input data [80, 88]. The decoding models then filter out the irrelevant noise as well as the inherent variations of the connectivity estimates [89]. We also equipped the model with squeeze-and-excitation blocks [90], which improved performance by enhancing the features of the most relevant frequency channels. On the contrary, the addition of spatial attention blocks [91] did not result in increased performance. Future work should consider further exploring attention strategies suitable for training deep networks with rs-FC input.

In this study, we focused on comparing models and evaluating input features instead of aiming for the highest possible predictive performance. Therefore, we decided to rely on cross-validation performance as a model comparison metric rather than utilizing a separate hold-out test set. Although applying a separate test set could reduce the risk of data leakage and consequent hyperparameter overfitting, it may also bias the testing error based on the random composition of the relatively small test set. By contrast, cross-validation allowed us to utilize all available data and rendered the performance metrics more robust to the limited sample size [92]. An inherent limitation of this approach is that cross validation, combined with early-stopping, may lead to optimistic predictive performance estimates. However, our previous results suggest that using a separate test set generally yields results comparable to cross-validation performance [93]. Furthermore, to ensure robust and comparable results, the 12-fold cross-validation procedure was repeated three times for each input and model with different random model initializations. Nonetheless, we cannot completely rule out the possibility of data leakage and overly optimistic results. Another limitation of the cross-validation approach is that the independence assumptions of the statistical tests applied are not necessarily fulfilled as the CV train–test splits overlap across folds. For this reason, the statistical results obtained through cross validation should be considered indicative only. Future work should consider including multiple independent datasets and verify that the proposed approach generalizes across sites and prediction targets.

We employed the chronological age of the participant as the prediction target for the decoding models. The difference between the predicted age and the chronological age is often termed the brain age delta. Although brain-predicted age may be useful as a biomarker in neurological and psychiatric disease [57, 58], the potential clinical utility of brain age prediction models has not been fully realized [86]. The use of brain age delta as a disease biomarker is further complicated by limited cross-site generalizability [94] and the often observed prediction bias towards the mean age of the particular dataset, which results in residual correlation between the brain age delta and chronological age unless explicitly corrected for [95]. Therefore, we encourage the development of direct neuroimaging biomarkers for specific pathologies, informed by normative modeling of the healthy population [10, 96]. In this study, we demonstrated the viability of our rs-FC decoding approach using the brain age prediction task. Next, it should be applied in the development of disease biomarkers.

A plethora of metrics to estimate functional connectivity from MEG and EEG recordings have been developed [97]. Unfortunately, different metrics do not always produce the same results, even on the same data, hindering the comparison of results across studies. Prior work has also reported variable test–retest reliability across metrics [76–78]. To avoid entering the swamp of endless connectivity measures, we limited the scope of this study to one representative metric from the two main families of amplitude-based and phase-based connectivity measures. Therefore, our results comparing AEC and wPLI do not necessarily generalize to other metrics in the respective families.

## 4 Conclusions

In this study, we proposed an interpretable deep learning framework for decoding brain states from MEG resting-state functional connectivity (rs-FC) data. We evaluated three proposed convolutional neural network (FC-CNN) models against standard machine learning approaches in the brain age prediction task and found that the proposed models outperformed the conventional regression methods. The best model produced a Mean Absolute Error (MAE) of 8.16 *±* 0.92 years, which is comparable to the state-of-the-art reported in other MEG-based brain age estimation studies [61–64]. We demonstrated how informative rs-FC patterns can be extracted from these models and showed that an ablated model with a small subset of the whole-brain rs-FC graph performed nearly as well as the complete model. We compared amplitude-based and phase-based rs-FC metrics and found that amplitude envelope correlation is more informative for decoding brain age than the weighted phase-lag index. Our results also suggest that to obtain optimal decoding performance, rs-FC should be estimated in an anatomical source space. These findings demonstrate that deep learning can be applied to extract useful information on brain states from functional connectivity data and will inform future development of neuroimaging-based biomarkers for brain disorders.

## 5 Methods

### 5.1 Participants and recordings

The data were collected at the Medical Research Council (UK) Cognition and Brain Sciences Unit (MRC-CBSU) in Cambridge, UK, as part of the Cambridge Center for Ageing and Neuroscience (Cam-CAN) study [45, 71]. The cross-sectional, multimodal open-access database includes structural and functional MRI, MEG, and behavioral recordings of a population-based cohort of nearly 700 healthy participants aged between 18 and 88 years with an approximately uniform age distribution. In this study, we included T1-weighted anatomical MRI and resting-state MEG data from *n* = 576 individuals. The duration of the resting-state MEG recording was 8 min 40 s. The primary study was conducted in accordance with the Declaration of Helsinki and was approved by the Cambridgeshire 2 Research Ethics Committee (reference: 10/H0308/50). The participants gave written informed consent. For a detailed description of the Cam-CAN study protocol, see the publication by Shafto and colleagues [71]. The data acquisition is detailed in the publication by Taylor and colleagues [45].

### 5.2 Preprocessing and functional connectivity analysis

The MEG data were processed using MNE–Python and MNE-Connectivity software tools [98–100]. The analysis is briefly described below and detailed in our previous publication [30].

#### MEG preprocessing

We employed spatiotemporal signal-space separation (tSSS; [101]) to suppress external interference and compensate for head movement during the measurement (MaxFilter v2.3; Megin Oy, Espoo, Finland). Subsequently, we filtered the data using a 1–100-Hz bandpass filter and a bandstop filter at the 50-Hz power line frequency and its harmonics. To suppress cardiac and oculomotor artifacts, we removed independent components displaying maximum cross-trial phase statistics with the ECG channel and maximum Pearson correlation with the EOG channels, respectively. We split the preprocessed data into 30-s epochs and removed any epochs containing residual artifacts. The subjects with at least 10 good epochs were retained for subsequent analyses.

#### Source estimation

We created individual cortical reconstructions from T1-weighted MRIs using FreeSurfer software [102, 103] and defined cortically-constrained source spaces comprising 5124 source points. We estimated source time series at these points using dynamic statistical parametric mapping (dSPM; [104]) and collapsed them into 448 cortical parcels following a parcellation developed by Khan and colleagues [23] based on the Desikan–Killiany atlas [105].

#### Functional connectivity analysis

We obtained analytic signals of parcel time series and sensor-level gradiometer signals by convolving them with a family of complex Morlet wavelets with 32 logarithmically-spaced central frequencies between 1 and 100 Hz. We estimated the degree of resting-state functional connectivity (rs-FC) between each pair of 448 parcels and between each pair of 204 planar gradiometers in each frequency bin using orthogonalized Amplitude Envelope Correlation (AEC; [21]) and weighted Phase-Lag Index (wPLI; [72]). We combined the resulting parcel connectivity estimates into 90 regions of interest (ROIs) based on the Desikan-–Killiany atlas [105] by averaging the connectivity estimates within each ROI [30]. The result is a set of *N*_nodes_ *× N*_nodes_ *× N*_freqs_ rs-FC estimates, where *N*_nodes_ = 90 and *N*_freqs_ = 32.

#### Normalization and combination of functional connectivity estimates

We applied min–max scaling to functional connectivity matrices for each subject and connectivity metric separately. Scaled AEC and wPLI estimates were concatenated into a shared connectivity tensor comrising the AEC estimates in the lower triangle and the wPLI estimates in the upper triangle for each frequency bin. This was done to probe if making phase and amplitude connectivity metrics available to the FC-CNN decoder will improve the predictive performance, as AEC and wPLI are known to contain non-redundant information [74].

### 5.3 Feature extraction

Most machine learning methods require a relatively small number of features to perform optimally. Although rs-FC matrix are already a compact representation of the MEG data, the total number of input features is still significantly larger than a typical number of training examples. We compared several dimensionality reduction techniques to alleviate overfitting which is a typical problem for datasets with low samples-to-features ratios. First, the functional connectivity estimates were averaged within the delta (1–4 Hz), theta (4–8 Hz), alpha (8–13 Hz), beta (13–30 Hz), and gamma (30–100 Hz) frequency bands. Second, we used principal component analysis (PCA) to combine 20025 individual source-space or 103530 sensor-level rs-FC estimates into approximately 400 principal components explaining 95% of the variance in the original data. PCA was fitted independently for each cross validation fold using only the training data.

Additionally, three graph-theoretical features (global efficiency, transitivity, and characteristic path length) were estimated using the density-based threshold corresponding to the top 20-th percentile for each frequency band separately. These graph metrics were chosen based on their demonstrated reproducibility on MEG data in previous studies [106]. The graph features were calculated using the NetworkX package [107].

### 5.4 Decoding models

We developed and evaluated three convolutional neural network (FC-CNN) architectures informed by assumptions about the statistical properties of the rs-FC data (Figure 2). FC-CNNs combine feature extraction and prediction in a single computational model while minimizing the number of trainable parameters to alleviate overfitting. We evaluated the performance of these models in the brain age prediction task. The models were built in MNEflow software [108]. We compared the performance of FC-CNNs to two machine learning models – support vector regression (SVR) and ridge regression (RR) – known for their successful performance with high-dimensional datasets [109, 110]. The models were fitted using Scikit-learn [111].

#### FC-CNN architectures

##### Input layer

A single input into FC-CNN, defined as a 3-dimensional tensor **X** *∈* ℝ*^N^*^nodes^*^×N^*^nodes^*^×N^*^freqs^, represents the functional connectivity estimates across all frequency bins. The first two dimensions represent the connectivity pattern of a given cortical parcel estimated at the frequency indexed along the third dimension.

##### Computational graph

The first layer of all proposed FC-CNN models extracts the multivariate connectivity pattern of each ROI by applying a set of global convolution kernels. We explored three variants of these connectivity pattern extraction layers. In the first variant, the *depthwise spatial convolution* model, the global spatial kernel **w_i_**is convolved with the input tensor

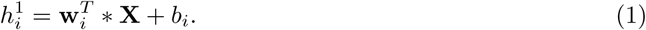

In the second variant, the *symmetric spatial convolution* model, the global spatial kernel **w_i_**is first convolved with the input tensor, and then the same kernel is convolved with the result to aggregate the weighted activity of nodes with similar first- and second-order proximity patterns.

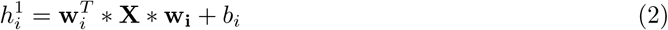

In both models above, spatial convolutions were performed for each frequency bin independently. This results in an *N*_nodes_ *× l × N*_freqs_ or *l × l × N*_freqs_ latent representation, where *l* is a free parameter defining the number of global convolution kernels. Separable convolutions are then followed by a 1 *×* 1 point-wise convolution along the frequency axis. To emphasize the most informative frequency bins, a squeeze-and-excitation channel-attention module (SE; [90]) is applied before the point-wise convolution layer.

By contrast, in the third variant, the *3D spatiospectral convolution* model, the model applies a 2-D global convolution across spatial and frequency dimensions. This is achieved by applying *l* two-dimensional convolution kernels with dimensions *N*_nodes_ *× N*_freqs_, followed by a 2-D global spatial convolution across the remaining spatial dimension. The advantage of this architecture is that it is capable of capturing connectivity patterns across frequency bins.

In each model, convolutional layers are followed by two fully-connected layers to obtain the brain age prediction result.

#### Training, cross-validation and model comparison

The training data consisted of frequency-resolved functional connectivity estimates from 576 individuals. We used mean absolute error (MAE) and *R*^2^-score estimated using 12-fold cross-validation for comparing predictive performance of different models and features. To compare model performance, we applied the two-sided Mann–Whitney U-test [112]. The results were corrected for multiple comparisons using Bonferroni correction.

For SVR and RR each training fold comprised an inner 5-fold cross-validation loop for optimizing regularization and kernel lengthscale parameters. Table 1 shows the hyperparameter search space.

**Table 1.** Hyperparameter grids for ridge regression (RR) and support vector regression (SVR) models.

| Model | Hyperparameter | Values |
| --- | --- | --- |
| RR | $\alpha$ | 50 values logarithmically spaced from $10^1$ to $10^4$ |
| SVR | $C$ | 10 values logarithmically spaced from $10^{-1}$ to $10^3$ |
| SVR | $\gamma$ | $\{10^{-4}, 5 \cdot 10^{-4}, 10^{-3}, 2.5 \cdot 10^{-3}, 5 \cdot 10^{-3}, 10^{-2}\}$ |

Each FC-CNN variant was trained using the Adam optimizer with early stopping after the validation MAE did not increase for 10 consecutive epochs. Each model was trained three times to obtain initialization-independent and comparable results. We evaluated several combinations of hyperparameters. Table 2 shows the evaluated parameter grid and the optimal combination.

**Table 2.**
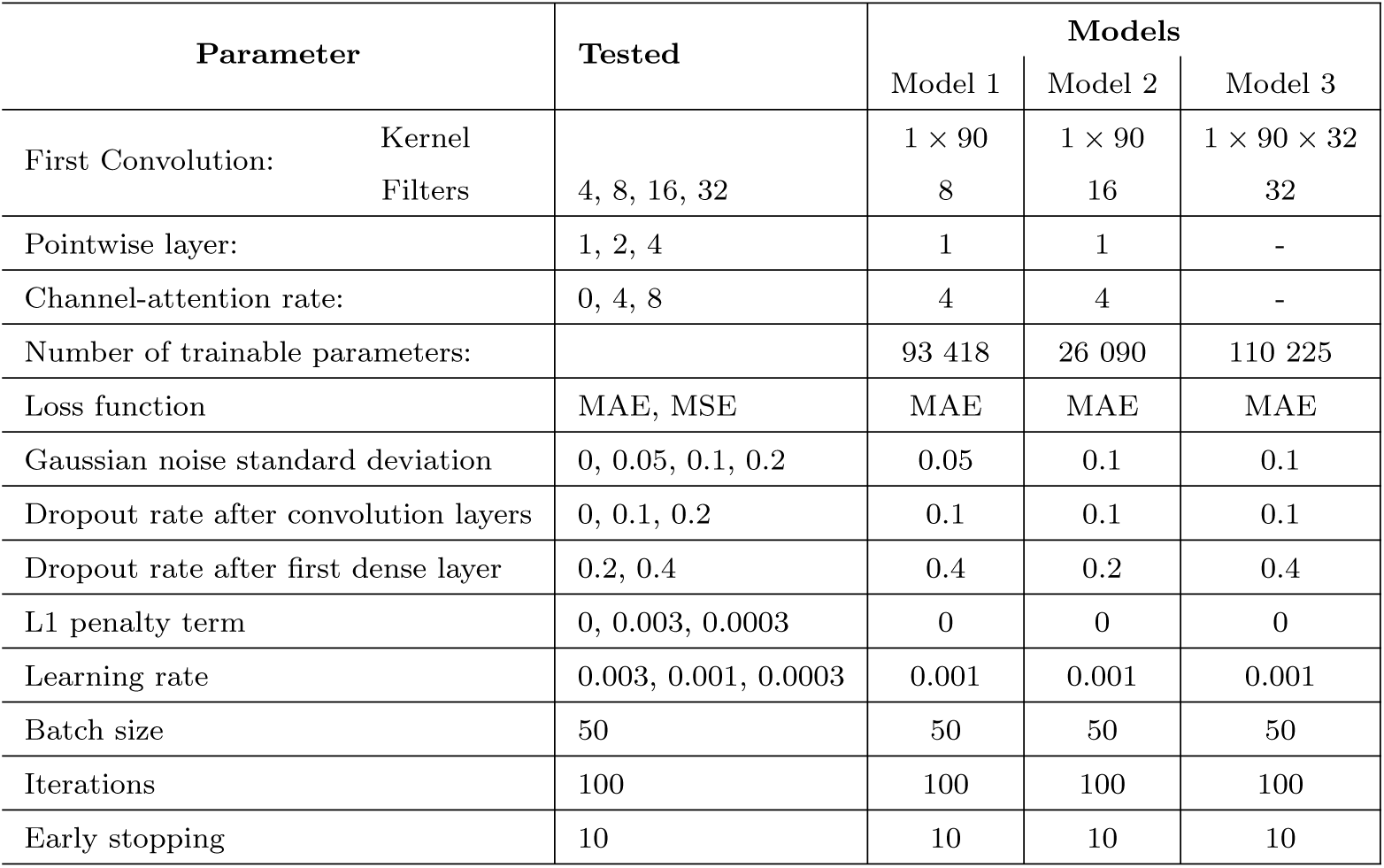
Parameters tested and selected in each layer of the FC-CNN models. The value 0 corresponds to exclusion of that layer from the final model.

**Table 3.** Evaluated sensor-level regularization levels.

| Parameter | Tested | Selected |
| --- | --- | --- |
| Gaussian noise standard deviation | 0.05, 0.1, 0.15 | 0.15 |
| Dropout rate after first dense layer | 0.2, 0.4 | 0.2 |

#### Augmentation and regularization

We employed several regularization methods to reduce overfitting. Most importantly, the network architecture was designed to have a minimal number of trainable parameters. Gaussian noise was injected into the connectivity matrices to prevent overdependence on minor fluctuations in the data. Gaussian noise is frequently used as an augmentation and regularization technique, for example, in image denoising [89]. In addition, we applied a sparse penalty on model weights, early stopping, dropout, and batch normalization techniques, summarized in Table 2.

### 5.5 Pattern interpretation and model ablation

FC-CNN learns complex multivariate features from the training data. The use of separable factorized convolutions allows grouping these features into simple interpretable embeddings. The suggested FC-CNN architectures combine global multivariate spatial and frequency-domain information into a single feature in the final prediction layer. Therefore, the feature with maximal weight in the output layer can be used to extract the corresponding spatial and frequency-domain activation patterns. Here, we derived activation patterns from the spatial convolution layers and used them to evaluate the subset of regions that contribute to successful age prediction. For each training fold, a single output-layer feature assigned the maximum weight was identified. Spatial pattern attributions were obtained following the procedure described by Kindermans and colleagues [113] using the spatial weights of the *symmetric depthwise convolution* model. The resulting activation patterns were normalized and averaged across folds. Next, we extracted ROIs corresponding to the top 20-th percentile of the grand-average activation pattern, and conducted an ablation study to confirm the contribution of the model-derived ROIs to age prediction. In the ablation study, we restricted input data to only the top 20% most active ROIs, retrained the model with the restricted input, and evaluated the ratio between the coefficients of determination of the ablated and full models 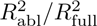.

## Data availability

The original MEG, MRI, and behavioral data are available upon request from the Cam-CAN data repository (https://camcan-archive.mrc-cbu.cam.ac.uk//dataaccess/).

## Code availability

The research code developed in this study is publicly available at https://github.com/ruuskas/dlfc-age-prediction. The FC-CNN models are available as part of *mneflow*, available at https://github.com/zubara/mneflow.

## Author contributions

S.R. and E.S. contributed equally to this study. According to the Contributor Role Taxonomy (CRediT), the authors contributed to the study as follows:

Conceptualization, Formal analysis, Investigation, Software – S.R., E.S., C.C., I.Z.

Data curation, Validation, Visualization – S.R.

Methodology – E.S., I.Z.

Writing – original draft – S.R., E.S., I.Z.

Writing – review & editing – S.R., E.S., C.C., L.P., I.Z.

Supervision, Project administration – L.P., I.Z.

Funding acquisition – L.P.

## Acknowledgments

This research was supported by the National Institute of Neurological Disorders and Stroke (award number 7R01NS104585-06 to Matti Hämäläinen) and Business Finland (grant “DIGIMIND” 7981/31/2022). Data collection and sharing for this project were provided by the Cambridge Center for Ageing and Neuroscience (Cam-CAN), funded by the UK Biotechnology and Biological Sciences Research Council (grant number BB/H008217/1), with support from the UK Medical Research Council and University of Cambridge, UK. We acknowledge the computational resources provided by the Aalto Science-IT project. We thank Matti Hämäläinen for providing resources for this study.

## Competing interests

The authors declare the following competing interests: L.P. has a part-time employment with the MEG device vendor Megin Oy. The other authors declare no competing interests.

## Notes

### Summary of Updates

Author list and author contributions updated to match the manuscript submitted to a journal.

